# Butterfly eggs prime anti-herbivore defense in an annual but not perennial *Arabidopsis* species

**DOI:** 10.1101/2024.06.05.597519

**Authors:** Maryse A. P. Huve, Norbert Bittner, Reinhard Kunze, Monika Hilker, Mitja N. P. Remus-Emsermann, Luis R. Paniagua Voirol, Vivien Lortzing

## Abstract

While plant anti-herbivore defenses of the annual plant species *Arabidopsis thaliana* were shown to be primable by *Pieris brassicae* eggs, the primability of the phylogenetically closely related perennial *Arabidopsis lyrata* has not yet been investigated. Previous studies revealed that closely related wild Brassicaceae plant species, the annual *Brassica nigra* and the perennial *Brassica oleracea*, exhibit an egg-primable defense trait, even though they have different life spans. Here, we tested whether *P. brassicae* eggs prime anti-herbivore defenses of the perennial *A. lyrata*. We exposed *A. lyrata* to *P. brassicae* eggs and larval feeding and assessed their primability by i) determining the biomass of *P. brassicae* larvae after feeding on plants with and without prior *P. brassicae* egg deposition and ii) investigating the plant transcriptomic response after egg deposition and/or larval feeding. For comparison, these studies were also conducted with *A. thaliana.* Consistent with previous findings, *A. thaliana*’s response to prior *P. brassicae* egg deposition negatively affected conspecific larvae feeding upon *A. thaliana*. However, this was not observed in *A. lyrata*. *Arabidopsis thaliana* responded to *P. brassicae* eggs with strong transcriptional reprogramming, whereas *A. lyrata* responses to eggs were negligible. In response to larval feeding, *A. lyrata* exhibited a greater transcriptome change compared to *A. thaliana*. Among the strongly feeding-induced *A. lyrata* genes were those that are egg-primed in feeding-induced *A. thaliana*, i.e., *CAX3, PR1*, *PR5* and *PDF1.4.* These results suggest that *A. lyrata* compensates for its lack of egg-mediated primability by a stronger response to larval feeding.

## Introduction

Plants employ various strategies to defend themselves against herbivorous insects, including constitutive and inducible anti-herbivore defenses. While constitutive defenses are always active, inducible anti-herbivore defenses are triggered only upon attack (Agrawal et al. 1999; Agrawal and Karban 1999; Karban and Baldwin 1997; Karban and Myers 1989; War et al. 2012). Inducibility may save energy, allowing resources to be allocated to other vital processes such as growth and reproduction (Cipollini et al. 2003). Initiating inducible defenses incurs a time delay, resulting in a time lag between an insect’s attack and the plant’s corresponding defense response. During this interval, the plant remains susceptible to herbivore damage (Frost et al. 2008). However, this delay can be reduced if the plant detects early cues indicating imminent insect attacks. This process, known as ‘priming’, prepares the plant’s defense mechanisms to “anticipate” and counter threats. As a result of priming, the plant’s response to herbivory becomes quicker, more sensitive, and potentially stronger, as it “anticipates” these attacks (Hilker et al. 2016; Hilker and Fatouros 2016; Hilker and Schmülling 2019; Martinez-Medina et al. 2016).

A wide range of environmental cues prime plant anti-herbivore defenses (Conrath et al. 2006; Frost et al. 2008; Pastor et al. 2013), for example, various volatile organic compounds, such as i) feeding damage-induced plant volatiles (Arimura et al. 2010; Dicke and Baldwin 2010; Kost and Heil 2006), ii) insect oviposition-induced plant volatiles (Pashalidou et al. 2020) or iii) insect pheromones (Bittner et al. 2019; Helms et al. 2013; Helms et al. 2014; Helms et al. 2017). Besides airborne cues, direct interactions with herbivorous insects, such as footprints (Peiffer et al. 2009), chewing vibrations (Appel and Cocroft 2014), and feeding damage (Rasmann et al. 2012) can prime plant anti-herbivore defenses. In addition, insect egg depositions on leaves are highly reliable cues indicating impending herbivory by hatching larvae (Hilker and Fatouros 2015, 2016).

Egg deposition by herbivorous insects was shown to prime a wide range of plant species, including tree species (Austel et al. 2016; Beyaert et al. 2012), perennial shrubs (Geuss et al. 2018; Pashalidou et al. 2015) and herbaceous annual species (Bandoly et al. 2015; Bandoly et al. 2016; Bonnet et al. 2017; Geiselhardt et al. 2013; Lortzing et al. 2019; Paniagua Voirol et al. 2020; Pashalidou et al. 2015; Rondoni et al. 2018). For annual plant species, egg primability is particularly beneficial because they produce seeds only once in their life cycle. Mitigation of severe damage by priming a plant’s anti-herbivore defense by prior insect egg deposition may support recovery from damage and thus, seed set (Valsamakis et al. 2022). However, also the anti-herbivore defense of perennial plants was shown to be primable by insect eggs (Austel et al. 2016; Beyaert et al. 2012; Geuss et al. 2018; Pashalidou et al. 2015; Pashalidou et al. 2020). Perennial plants might have long term benefits from priming their anti-herbivore defenses not only for the current season, but also the future one (Haukioja et al. 1985; Schott et al. 2023). The evolution of primable traits in annual and perennial plants might be driven by various factors, among them the plant’s “memory” abilities as well as environmental factors like the predictability of stress or the community in which the organism is living (Hilker et al. 2016).

Previous studies highlighted that egg deposition by the Large White butterfly (*Pieris brassicae*) primes various Brassicaceae species, leading to an impaired development of *P. brassicae* larvae. *Pieris brassicae* larvae feeding on prior egg-laden annual *Arabidopsis thaliana* consumed less leaf tissue, gained less weight and suffered higher mortality in comparison to *P. brassicae* larvae feeding on egg-free plants (Geiselhardt et al. 2013; Valsamakis et al. 2022). This effect is elicited by a secretion attached to *P. brassicae* eggs (Paniagua Voirol et al. 2020) and is mediated by the phytohormone salicylic acid (SA) (Lortzing et al. 2019). Furthermore, egg-primed responses are adjusted to the hatching time point of *P. brassicae* larvae (Valsamakis et al. 2020). Apart from *A. thaliana*, also the defenses of other annual Brassicaceae are primable by *P. brassicae* eggs: *Brassica nigra*, *Sinapis arvensis* and *Moricandia moricandioides* (Pashalidou et al. 2013; Pashalidou et al. 2015). So far, the primability of anti-herbivore defenses by *P. brassicae* eggs has only been described in one perennial Brassicaceae, *Brassica oleracea* (Pashalidou et al. 2015; Pashalidou et al. 2020). This raises the question whether also other perennial Brassicaceae species are primable by butterfly eggs and whether their primability is comparable to annual Brassicaceae species.

The aim of this study was to investigate the egg primability of the perennial *Arabidopsis lyrata* (Koch et al. 1999; Price et al. 1994) and to compare it with the well-known egg primability of the phylogenetically closely related, annual species *A. thaliana* (Geiselhardt et al. 2013; Lortzing et al. 2019; Paniagua Voirol et al. 2020; Valsamakis et al. 2020; Valsamakis et al. 2022). Both plant species share a high degree of genome sequence similarity (Koch et al. 1999; Nasrallah 2000), but have different life spans and grow in different environmental conditions (Al-Shehbaz and O’Kane 2002). *Arabidopsis thaliana* occurs as a winter annual on sandy soil, roadsides, rocky slopes, waste places, cultivated ground and meadows (Al-Shehbaz and O’Kane 2002), whereas *A. lyrata* is a drought-tolerant pioneer herb occurring in low-competition environments such as cliffs, calcareous ledges, rock crevices and sandy areas, e.g., with calcium-deficient serpentine soils (Al-Shehbaz and O’Kane 2002; Clauss and Koch 2006; Koenig and Weigel 2015; Mitchell-Olds 2001; Nasrallah 2000; Sletvold and Ågren 2012; Turner et al. 2010). Based on these differences, we asked whether *A. thaliana* and *A. lyrata* respond differently to insect egg deposition.

A previous study showed that *P. brassicae* eggs induce an egg-killing trait that is phylogenetically conserved within species of the Brassiceae tribe including *Brassica* crops and close relatives (Griese et al. 2021). Furthermore, the anti-herbivore defenses of the closely related *B. nigra* and *B. oleracea* plant species are primable by *P. brassicae* eggs (Pashalidou et al. 2013; Pashalidou et al. 2015; Pashalidou et al. 2020), although they have different life spans. We hypothesized that the anti-herbivore defenses of *A. lyrata* are as primable by *P. brassicae* eggs as those of *A. thaliana*. Therefore, we compared the primability of *A. thaliana* and *A. lyrata* by exposure of both plant species to *P. brassicae* eggs and larval feeding. We i) determined the biomass of larvae after feeding on plants with and without prior egg deposition and ii) investigated the plant transcriptomic response after egg deposition and/or larval feeding. Unlike *A. thaliana*, defenses of *A. lyrata* against *P. brassicae* larval feeding are not primable by *P. brassicae* eggs. Instead, *A. lyrata* exhibited a greater transcriptome change in response to larval feeding compared to *A. thaliana*. These results suggest that *A. lyrata* compensates for its lack of egg-mediated primability by a stronger response to larval feeding.

## Material and methods

### Plant material and growth conditions

Seeds of *Arabidopsis thaliana* (Columbia-0) and *A. lyrata* ssp. *lyrata* (RonC) were sown on a 3:1 mixture of soil (Einheitserde classic):sand and stratified for two days at 4 °C. The plants grew in climate chambers under short day conditions (10 h/ 14 h light dark cycle, 21 °C, 40% relative humidity, 100-120 µmol m^-2^ s^-1^ light intensity).

### Pieris brassicae rearing

*Pieris brassicae* was reared as described by Valsamakis et al. (2022). Briefly, larvae and adult butterflies were kept in flight cages (45 cm x 45 cm x 60 cm) under long day conditions (18 h/6 h light/dark cycle, 220 µmol m^-2^ s^-1^ light intensity, 23 °C and 70% relative humidity). The larvae were fed with Brussels sprouts plants (*Brassica oleracea* var. *gemmifera*) until pupation. The adult butterflies were fed with 15% aqueous honey solution. Mated female butterflies were regularly offered Brussels sprouts plants for egg deposition.

### Plant treatments

#### Treatment with eggs

We treated *Arabidopsis* plants by exposing a single fully developed, non-senescent leaf (leaf position 17-22) to one mated female butterfly, resulting in the deposition of a clutch containing 30-40 *P. brassicae* eggs. After six days under short day conditions (10 h/14 h light-dark cycle, 21 °C, 40% relative humidity, 100-120 µmol m^-2^ s^-1^ light intensity), we carefully removed the eggs using tweezers and a fine brush. Egg-free plants were used as control.

#### Treatment with larvae

One day before hatching, *P. brassicae* eggs deposited on Brussels sprouts were collected and placed in Petri dishes until the larvae hatched (10 h/14 h light-dark cycle, 21 °C, 40% relative humidity, 100-120 µmol m^-2^ s^-1^ light intensity). A group of ten neonate larvae was placed on a single, fully developed, non-senescent leaf (leaf position 17-22) of a previously egg-laden or an egg-free *A. thaliana and A. lyrata* plant, respectively, and enclosed in a clip cage (2 cm in diameter, 1.7 cm high). After two days feeding within the clip cage, larvae were removed from the plant for biomass determination (see below). Thereafter, they were placed back to their respective plants and were allowed to freely move and feed on the entire plant. To prevent larvae from escaping, the plants were enclosed in Plexiglas® cylinders (14.5 cm diameter, 15 cm high) with a gauze lid.

### Determination of larval biomass

Larval performance was assessed by determining the biomass of *P. brassicae* larvae per plant with a fine-scale balance (Ohaus® Analytical Plus balance Ohaus AP250D, Nänikon, Schweiz). We calculated the average larval biomass after a two-day and five-day feeding period on seven-week-old *A. thaliana* and *A. lyrata* plants. Additionally, the larval biomass was determined after a two-day and five-day feeding period on nine-week-old, egg-free and previously egg-laden *A. lyrata* plants.

### Sampling of leaf tissue

For transcript analyses, the experiments were designed in 2 x 2 factorial setup. Plants were exposed to eggs (E), to larval feeding (F), or to both eggs and feeding (E+F). Untreated plants were used as control (C). After two days of feeding on seven-week-old plants with or without eggs, we harvested leaf material for transcriptome analyses. Treated leaves were flash frozen in liquid nitrogen.

### RNA extraction

We extracted total RNA as described by Oñate-Sánchez and Vicente-Carbajosa (2008) and removed residual genomic DNA with the TURBO DNA free™ kit (ThermoFisher Scientific, Waltham, USA) following the manufacturers recommendations. The RNA quantity and quality was inspected on a 1.2% agarose gel and with a Multiscan^®^ GO Microplate Spectrophotometer (Thermo Scientific). RNA integrity (RIN between 6.6-8.8) was estimated using the Bioanalyzer 2100 (Agilent Technologies, Santa Clara, USA) before samples were sequenced (Macrogen, Europe).

### RNA sequencing and analysis of differentially expressed genes

The lllumina TruSeq Stranded Poly-A selected RNA Sample library kit for plants was used to prepare samples for sequencing. Paired end sequencing (2 x 150 bp) was conducted using the NovaSeq6000 platform (Illumina, San Diego, USA). All samples produced between 32.5 and 36.6 million reads. For adapter clipping and trimming Trimmomatic was used (version 0.39) (Bolger et al. 2014). Sequences shorter than 50 bp were excluded from further analysis. The sequence quality was inspected with FastQC and MultiQC before and after adapter clipping and trimming (Andrews 2020; Ewels et al. 2016). Ribosomal sequences were filtered with SortmeRNA (version 2.1) (Kopylova et al. 2012). To map reads against the plant genomes, the genomes and their annotation from *A. thaliana* and *A. lyrata* were obtained from Ensembl Plants (Howe et al. 2020) (version TAIR10, release 44) and from Rawat et al. (2015). Reads were counted with kallisto (version 0.46.0) (Bray et al. 2016), and resulting count files were converted to the DESeq2 package data format with the tximeta package (Love et al. 2020) (Bioconductor version 3.9) (Soneson et al. 2015) in R (R Core Team 2022). All genes with a read count > 1 were considered for analysis of differential expression. Differentially expressed genes (DEGs) were defined to have a *P* value ≤ 0.05 after *fdr* correction for multiple testing (Benjamini and Hochberg 1995).

For further downstream analysis of gene functions, we annotated *A. lyrata* genes to their *A. thaliana* orthologs by employing supplemental datasets of the latest *A. lyrata* annotation (S4 dataset) (Rawat et al. 2015). The biological functions of differentially expressed genes (DEGs) were examined via enrichment analyses of Kyoto Encyclopedia of Genes and Genomes (KEGG) pathway and gene ontology (GO) terms with DAVID 6.8 (Da Huang et al. 2009; Sherman et al. 2022). The transcriptome data were further explored using R (R Core Team 2022) integrated in RStudio (version 2022.12.0) (RStudio Team 2022) and venn diagrams [package “eulerr” (Larsson 2024)], hierarchical clustering analyses, heatmapping using Euclidean distances, and Complete-linkage clustering [packages “ComplexHeatmap” (Gu et al. 2016), “dentextend” (Galili 2015)].

### cDNA synthesis and quantitative real-time PCR

First strand cDNA was synthesized from 2 µg RNA with the smART Reverse Transcriptase kit (Roboklon GmbH, Berlin, Deutschland) and oligo-dT18 following the manufactureŕs protocol. Quantitative real-time PCRs were conducted in a total of 10 µl using Blue ŚGreen qPCR 2x Mix (Biozym Scientific GmbH, Hessisch Oldendorf, Deutschland). As reference genes, for *A. thaliana ACT2* (AT3G18780), *GADPH* (AT1G13440) and *TUB6* (AT5G12250), and for *A. lyrata PS2SUB, PYRT* and *PPP2R1’3* were used. All primers are listed in Supplementary Table S1. Relative expression of each gene was calculated with the ΔΔCT method (Livak and Schmittgen 2001).

### Statistics

Statistical evaluation and visualizing were performed with R (R Core Team 2022; RStudio Team 2022). The following packages were used: car (Fox and Weisberg 2019), cowplot (Wilke 2022), ggplot2 (Wickham 2016), ggpubr (Kassambara 2023), psych (Revelle 2024), reshape2 (Wickham 2007), Rmisc (Hope 2022), tidyverse version 1.3.0 (Wickham et al. 2019), viridis (Garnier et al. 2024).

Data distribution was evaluated with Shapiro-Wilk test and Q-Q-plot. Homogeneity of data variances was assessed using Levene’s test. For larval biomass data, we applied multiple Student’s *t*-test with *fdr* correction *post hoc* (Benjamini and Hochberg 1995), because data were normally distributed and had homogenous variances. For qPCR data, we applied 2×2 factorial ANOVA with Tukey *post hoc* test based on normal distribution of data with homogenous variances.

## Results

### The plant’s response to *Pieris brassicae* eggs negatively affects larval growth on *Arabidopsis thaliana* but not on *Arabidopsis lyrata*

We assessed larval biomass after two and five days of feeding on seven-week-old *A. thaliana* and *A. lyrata*, with or without prior exposure to *P. brassicae* eggs (Fig. 1). Larvae feeding on previously egg-laden *A. thaliana* plants gained significantly less weight compared to those feeding on egg-free plants, regardless of the feeding duration. By contrast, prior egg deposition on *A. lyrata* did not affect larval biomass. This lack of an egg-priming effect on larval biomass was observed in both seven-week-old (Fig. 1) and nine-week-old *A. lyrata* plants (Fig. S1). Moreover, larvae gained significantly more biomass when feeding on *A. lyrata* compared to *A. thaliana* (Fig. 1).

**Fig. 1.**
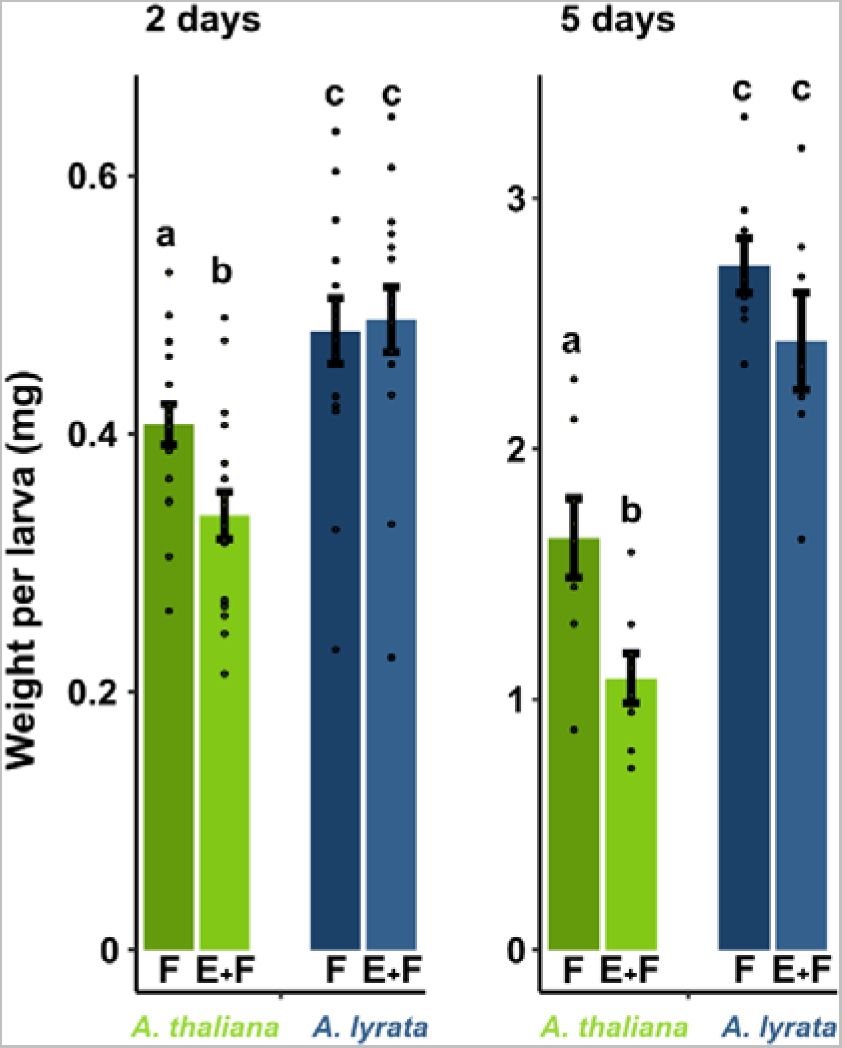
Impact of the plant’s responses to *Pieris brassicae* eggs on biomass of conspecific larvae feeding on *Arabidopsis thaliana* (green) and *A. lyrata* (blue) plants. Biomass in mg (means ± SE) of larvae after 2 or 5 days feeding on previously egg-laden (E+F) and egg-free (F), seven-week-old plants. Dots represent the data points. Different letters above the bars indicate significant differences between treatments and plant species (*P* < 0.05, multiple Student’s *t*-test with *fdr* correction). After 2 days feeding *N* = 16-18, after 5 days feeding *N* = 7-8. Statistical details are provided in Supplementary Table S2

### *Arabidopsis lyrata* responds to *Pieris brassicae* eggs with a weaker transcriptional reprogramming than *Arabidopsis thaliana*

We investigated the effect of *P. brassicae* eggs on the transcriptomes of seven-week-old *A. lyrata* and *A. thaliana* plants using RNA-seq. The eggs remained on the plant leaves for six days, which is the typical incubation time until hatching at 20 °C (David and Gardiner 1962) and were removed just before hatching. In response to *P. brassicae* eggs, *A. lyrata* showed fewer differentially expressed genes (DEGs) than *A. thaliana* (Fig. 2). *Arabidopsis thaliana* responded to *P. brassicae* eggs with strong transcriptional reprogramming. Overall, 1396 *A. thaliana* genes were differentially expressed (1002 up-, 394 downregulated genes) (Fig. 2, Supplementary Data 1). This represents 5% of the total number of protein coding genes of *A. thaliana* (Cheng et al. 2017). The transcriptomic response of *A. lyrata* to *P. brassicae* eggs was much weaker compared to *A. thaliana*, with *A. lyrata* exhibiting 326 upregulated genes and 19 downregulated genes in response to the eggs (Fig. 3, E vs C, Supplementary Data 1), which represents 1% of the total number of protein coding genes in *A. lyrata* (Hu et al. 2011).

**Fig. 2.**
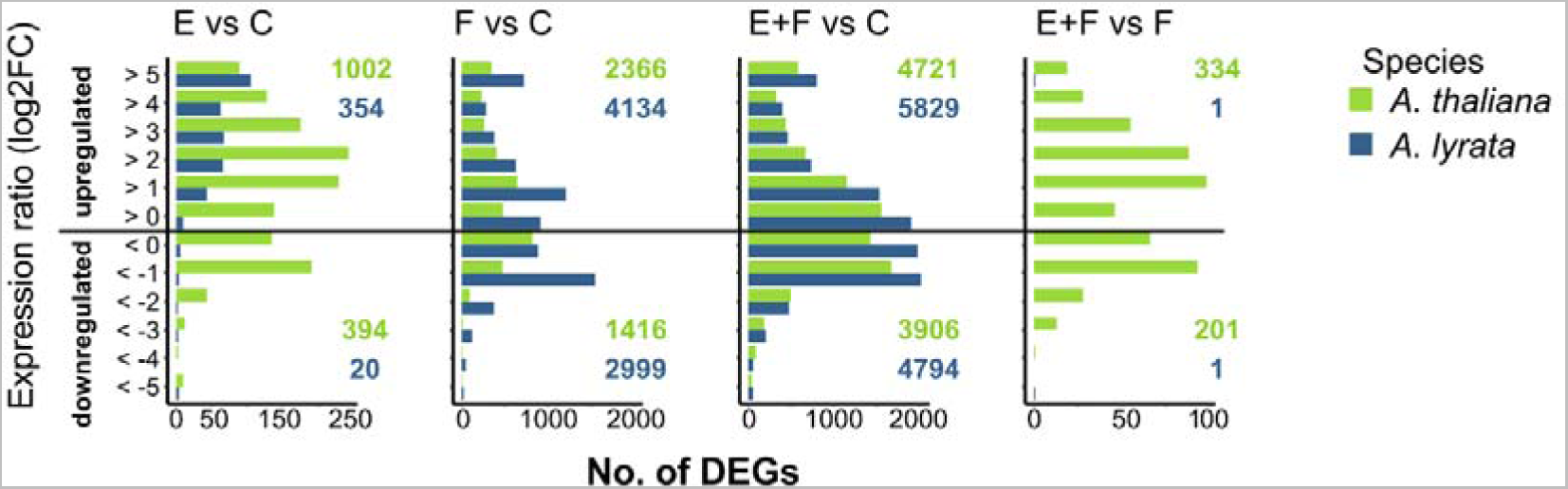
Number of differentially expressed genes (DEGs) in *A. thaliana* (green) and *A. lyrata* (blue) in response to *Pieris brassicae* eggs (E), larval feeding (F) or both, eggs followed by larval feeding (E+F) for the following treatment comparisons: E versus C, F versus C, E+F versus C and E+F versus F. Control plants (C) were left untreated. *N* = 5.

**Fig. 3.**
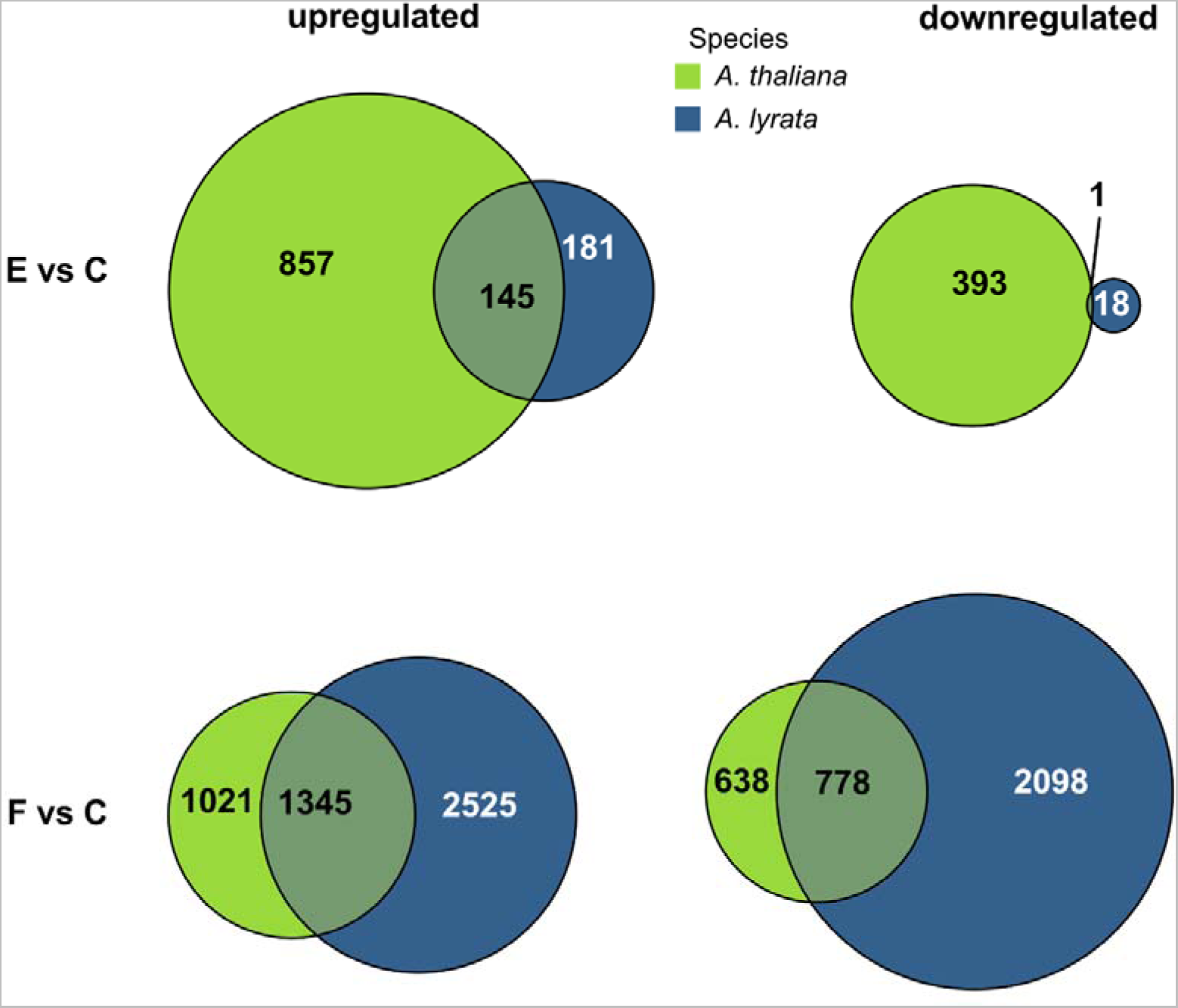
Common and unique differentially expressed genes in *Arabidopsis thaliana* and *A. lyrata* in response to *Pieris brassicae* eggs (E vs C) or larval feeding (F vs C). Circular areas are drawn to scale relative to the number of DEGs. *Arabidopsis lyrata* genes without *A. thaliana* orthologs are not shown in this figure but are listed in Supplementary Data 1. *N* = 5

In the following, we will analyze the transcriptomic data sets with respect to the plant responses to eggs, to larval feeding and to the sequence of egg laying and larval feeding. When differentiating between up- and downregulated DEGs, we will especially focus on genes involved in phytohormone signaling [jasmonic acid (JA), SA, abscisic acid (ABA)], in secondary metabolite biosynthesis (phenylpropanoids, glucosinolates), photosynthesis, calcium and oxidative stress signaling, plant-pathogen interactions [e.g., systemic acquired resistance (SAR), and unfolded protein responses]. We focus on these gene biological functions because they are known to play a role in plant responses to insect egg deposition, insect feeding, or both (Geuss et al. 2018; Little et al. 2007; Lortzing et al. 2017; Lortzing et al. 2020; Valsamakis et al. 2022).

#### Upregulation of genes in response to oviposition

Nearly half (44%) of the orthologous egg-inducible genes in *A. lyrata* were also found to be inducible in *A. thaliana* (Fig. 3, E vs C, Supplementary Data 4). These egg-inducible genes in both *Arabidopsis* species include well-known insect egg-responsive markers like *PR5* and *PDF1.4*, which showed similarly strong egg inducibility in both plant species (Supplementary Data 4). In an independent experiment, qPCR analysis confirmed that both plant species exhibited similarly strong expression of *PDF1.4, PR1* and *PR5* in response to eggs, while the expression of *CAX3* was considerably stronger in *A. thaliana* than in *A. lyrata* (Fig. 4).

**Fig. 4.**
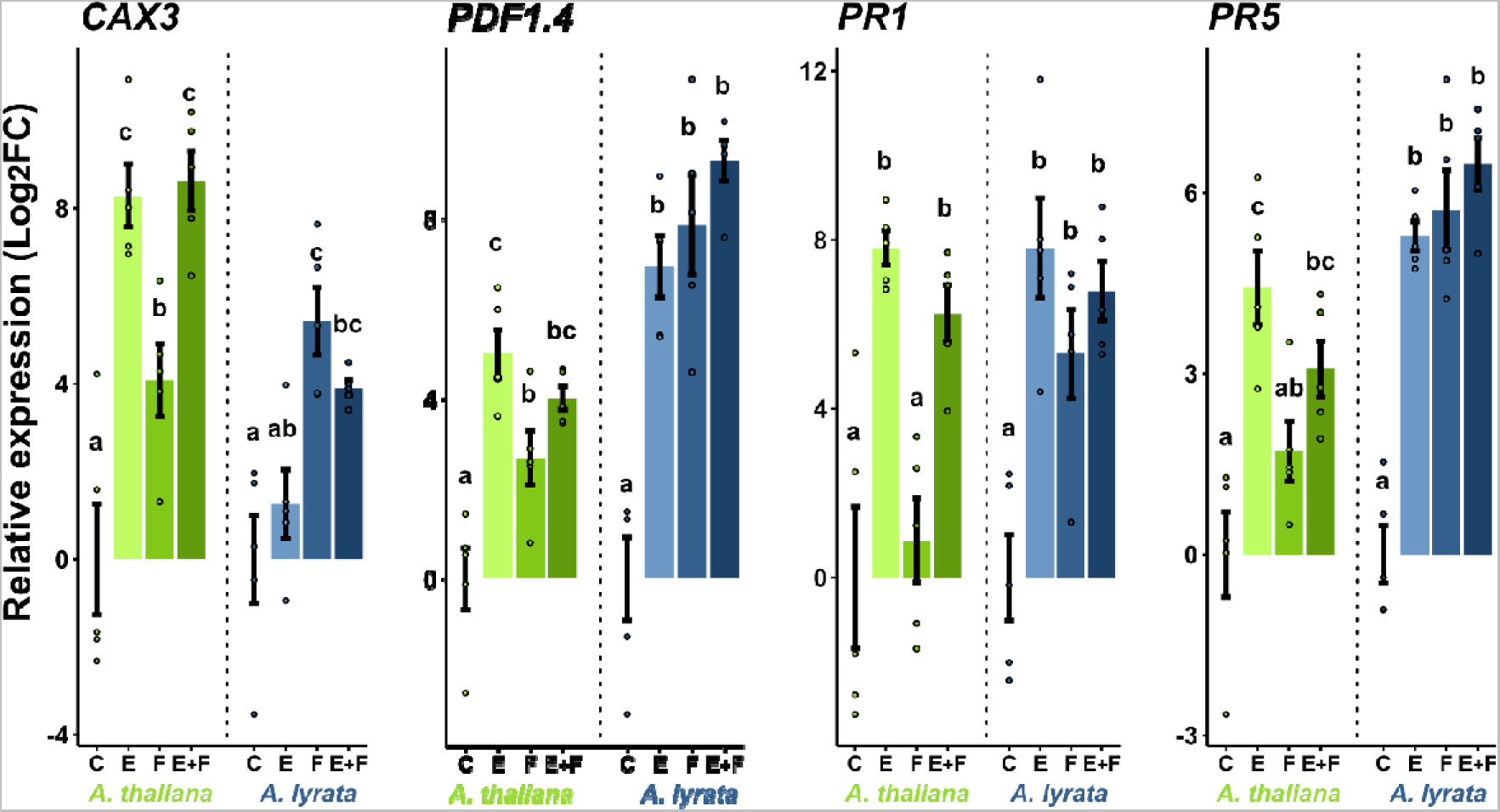
Relative expression of the priming-responsive genes *CAX3*, *PDF1.4*, *PR1* and *PR5* in *Arabidopsis thaliana* (green) and *A. lyrata* (blue). The plants were exposed to *Pieris brassicae* eggs (E), larval feeding (F) or eggs with subsequent larval feeding (E+F) or were left untreated (C). Bars indicate mean relative expression (Log_2_FC) ± SE, dots represent individual data points. Different letters above the bars indicate significant differences between treatments (*P* < 0.05, 2×2 ANOVA with Tukey test *post hoc*). Detailed statistics are provided in Supplementary Table S3. *N* = 4-5

The upregulated genes in egg-laden *A. lyrata* were enriched in less KEGG pathway and GO terms than in *A. thaliana* [KEGG pathway term enrichment: 6 terms for *A. lyrata* and 10 terms for *A. thaliana* (Fig. 5, Supplementary Data 2); GO term enrichment: 63 terms for *A. lyrata* and 98 terms for *A. thaliana* (Supplementary Data 3)].

**Fig. 5.**
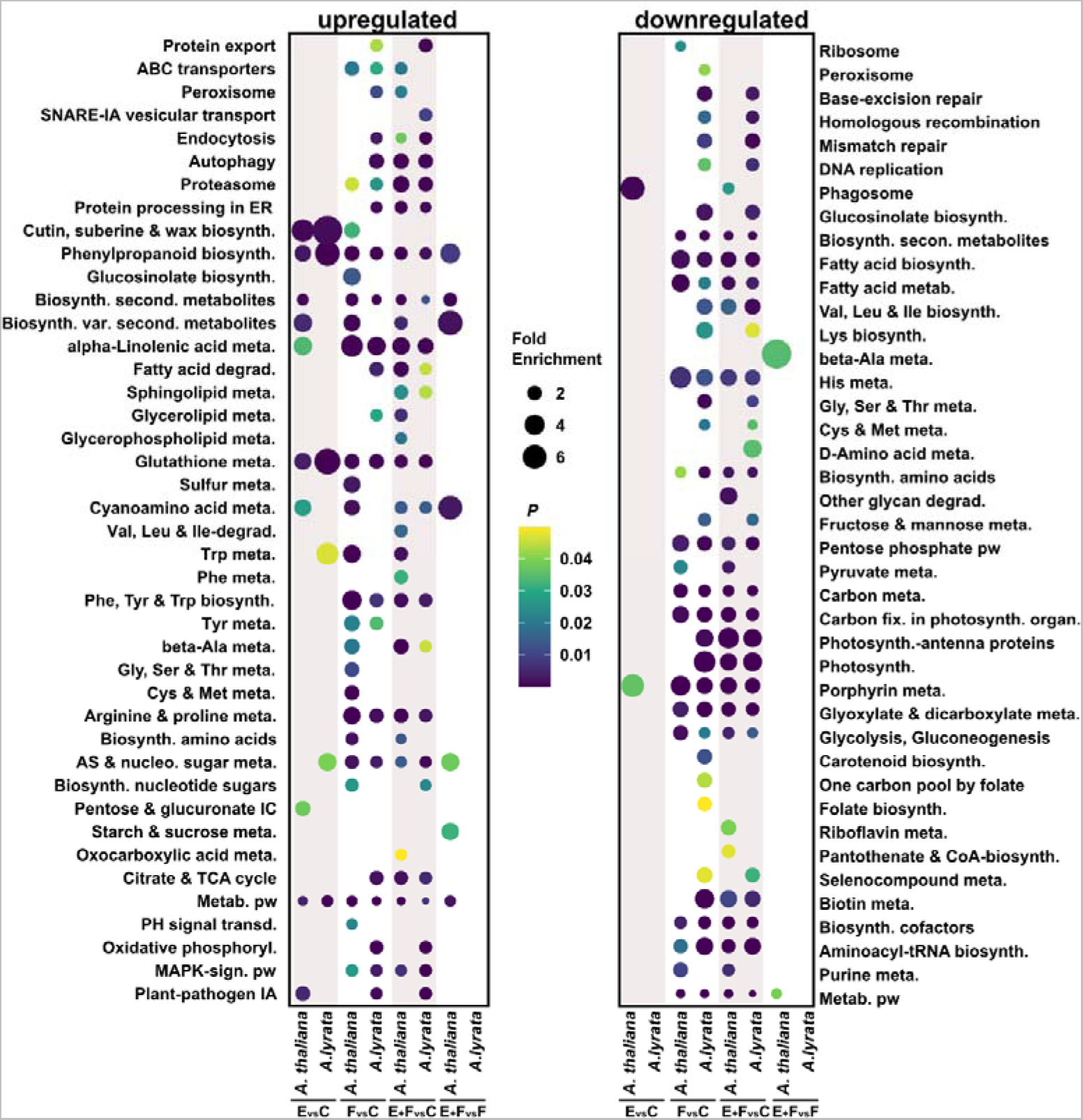
Transcriptional responses of *Arabidopsis thaliana* and *A. lyrata* to *Pieris brassicae* eggs, larval feeding or both, eggs followed by larval feeding. Seven-week-old plants were exposed to *P. brassicae* eggs (E), larval feeding (F), eggs and larval feeding (E+F) or were left untreated (C). a) Number of differentially expressed genes (DEGs) in *A. thaliana* (green) and *A. lyrata* (blue) and b) their enrichments in KEGG pathways for the following treatment comparisons: E versus C, F versus C, E+F versus C and E+F versus F. For b) the circle size represents fold enrichment of genes in the KEGG pathway, the color indicates the *P* value. *N* = 5. Abbreviations: Ala: alanine, AS: amino sugar, biosynth.: biosynthesis, Cys: cysteine, degrad.: degradation, ER: endoplasmic reticulum, fix: fixation, Gly: glycine, His: histidine, IA: interaction, IC: interconversions, Ile: isoleucine, Leu: leucine, Lys: lysine, Met: methionine, meta.: metabolism, metab.: metabolic, nucleo: nucleotide, PH: plant hormone, Phe: phenylalanine, phosphoryl.: phosphorylation, pw: pathway, second.: secondary, Ser: serine, sign.: signaling, TCA: tricarboxylic acid, Thr: threonine, transd.: transduction, Trp: tryptophan, Tyr: tyrosine, Val: valine, var.: various

Similar to *A. thaliana*, upregulated genes in *A. lyrata* in response to *P. brassicae* eggs were enriched in the KEGG pathway term phenylpropanoid biosynthesis [e.g., *PEROXIDASE 52* (*PRX52*)] as well as in GO terms related to SAR (e.g., *PR5*), SA-mediated signaling, e.g., *WRKY DNA-BINDING PROTEIN*s including *WRKY18, WRKY46, WRKY60*), JA-mediated signaling [e.g., *JASMONIC ACID OXIDASE* 3 (*JOX3*)], and oxidative stress [e.g., *SENESCENCE ASSOCIATED GENE 14* (*SAG14*)] (Fig. 5, Supplementary Data 2 and 3).

However, in *A. lyrata* fewer genes were enriched in these GO terms than in *A. thaliana*. For example, specific genes like *DIHYDROFLAVONOL 4-REDUCTASE* (*TT3*), *ANTHOCYANIDIN SYNTHASE* (*ANS*), *FLAVONOL SYNTHASE* (*FLS*) *5* (phenylpropanoid biosynthesis), *ALLENE OXIDE CYCLASE* (*AOC*) *1*, *AOC3*, and *OXOPHYTODIENOATE-REDUCTASE 3* (*OPR3*), *JASMONATE INSENSITIVE 1* (*MYC2*), (JA pathway), *AVRPPHB SUSCEPTIBLE 3* (*PBS3*), *ENHANCED DISEASE SUSCEPTIBILITY 5* (*EDS5*)] (SA pathway), were upregulated only in previously egg-laden *A. thaliana* (Supplementary Data 3 and Data 4). Furthermore, *A. thaliana* responded to eggs with the upregulation of genes enriched in GO terms associated to calcium-mediated signaling and calcium ion homeostasis, e.g., 8 *CALMODULIN-LIKE* (*CML*) genes, *CALMODULIN 3* (*CAM3*) and *CALMODULIN 8* (*CAM8*), 3 *CALCIUM-DEPENDENT PROTEIN KINASE* (*CPK*) genes and *CAX3* (Supplementary Data 3 and 4). This response to *P. brassicae* eggs is absent in *A. lyrata* (Supplementary Data 3). This is also reflected by our qPCR data showing that *CAX3*, encoding for a Ca^2+^/H^+^ exchanger (Manohar et al. 2011), is upregulated in egg-laden *A. thaliana*, but not in egg-laden *A. lyrata* (Fig. 4).

#### Downregulation of genes in response to oviposition

*Arabidopsis lyrata* downregulated only 20 genes in response to *P. brassicae* eggs (less than 0.5% of all downregulated genes in *A. lyrata*), whereas *A. thaliana* downregulated 394 genes in response to *P. brassicae* eggs (43% of all downregulated genes in *A. thaliana*) (Fig. 2).

Due to the limited number of downregulated genes in *A. lyrata* in response to eggs, the KEGG and GO term analyses lacked sufficient counts for meaningful interpretation. Genes downregulated in egg-laden *A. thaliana* were enriched in KEGG pathways and GO terms linked to chlorophyll biosynthesis [e.g., porphyrin metabolism, *PROTOCHLOROPHYLLIDE OXIDOREDUCTASE A* (*PORA*)], cell cycle and cell division [e.g., *CELL DIVISION CYCLE 20.1* (*CDC20.1*)] (Fig. 5, Supplementary Data 2 and Data 3).

Only one gene, *GDPDL4*, was downregulated in both *A. lyrata* and *A. thaliana* in response to eggs (Fig. 3, E vs C, Supplementary Data 4). *GDPDL4* encodes a protein with glycerophosphoryl diester phosphodiesterase-like activity, which plays a role in cellulose accumulation and pectin linking (Hayashi et al., 2008).

### *Arabidopsis lyrata*’s transcriptome response to *Pieris brassicae* larval feeding is stronger than that of *A. thaliana*

We determined the impact of *P. brassicae* larval feeding on the transcriptomes of seven-week-old *A. lyrata* and *A. thaliana* plants. Plants were exposed to larval feeding for a period of two days.

Both *A. lyrata* and *A. thaliana* exhibited a strong transcriptional reprogramming in response to larval feeding (Fig. 2, Supplementary Data 1). The transcriptional reprogramming in *A. thaliana* was stronger, when plants were previously exposed to the eggs (Fig. 2, E+F vs. F). This could not be observed in *A. lyrata*. However, upon feeding damage alone (F vs C), 1.6 times more genes were up- or down-regulated in *A. lyrata* than in *A. thaliana*, with 7142 protein coding genes showing a response in *A. lyrata* (4143 up- and 2999 down-regulated genes) compared to 3782 genes in *A. thaliana* (2366 up- and 1416 down-regulated genes) (Fig. 2, Supplementary Data 1).

When considering the genes with orthologs in both species, more than 50% that were regulated in *A. thaliana* by larval feeding were also regulated in *A. lyrata* (57% for upregulated, 55% for downregulated genes). However, *A. lyrata*’s transcriptional reprogramming was stronger in response to larval feeding than that of *A. thaliana* (Fig. 3, F vs C, Supplementary Data 4). The qPCR-analyses of egg-priming responsive genes also indicated that *CAX3*, *PDF1.4*, *PR1* and *PR5* were much stronger induced in *A. lyrata* in response to larval feeding than in *A. thaliana* compared to the untreated control plants (Fig. 4).

#### Upregulation of genes in response to larval feeding

Although larval feeding induced more of the orthologous genes in *A. lyrata* than in *A. thaliana*, the KEGG pathway and GO term analysis resulted in similar numbers of enriched terms for both plant species [KEGG pathway term enrichment: 22 terms for *A. lyrata* and 24 terms for *A. thaliana* (Fig. 5 and Supplementary Data 2); GO term enrichment: 166 terms for both plant species (Supplementary Data 3)].

Both in *A. lyrata* and in *A. thaliana*, upregulated genes responding to larval feeding were significantly enriched in KEGG pathway terms associated with phenylpropanoid biosynthesis, biosynthesis of secondary metabolites, glutathione metabolism and alpha-linolenic acid metabolism (Fig. 5). Furthermore, commonly upregulated genes were enriched in GO terms associated to ABA [*RESPONSE TO ABA AND SALT 1* (*RAS1*), *NDR1/HIN1-LIKE 6* (*NHL6*)], JA [*ACYL-COA OXIDASE 1* (*ACX1*), *LIPOXYGENASE 2* and *3* (*LOX2, LOX3*)] and SA synthesis and signaling *ENHANCED DISEASE SUSCEPTIBILITY* 5 (*EDS5*), *12-OXOPHYTODIENOATE REDUCTASE 1* (*OPR1*)], and to the synthesis of flavonoids [*TRANSPARENT TESTA 8* (*TT8*)], including anthocyanins [*ANTHOCYANIDIN SYNTHASE* (*ANS*), *PRODUCTION OF ANTHOCYANIN PIGMENT* (*PAP1*)] (Supplementary Data 3 and Data 4).

Additionally, in *A. lyrata*, the feeding-induced genes were significantly enriched in a KEGG pathway term involved in plant-pathogen interactions (e.g., *CPK32*, *CPK1*) (Fig. 5, Supplementary Data 2). In comparison to *A. thaliana*, more feeding-induced *A. lyrata* genes were enriched in GO terms associated to unfolded protein response (e.g., endoplasmic reticulum unfolded protein response and response to endoplasmic reticulum stress) and to SA signaling (e.g., regulation of salicylic acid biosynthetic process and regulation of salicylic acid mediated signaling pathway) (Supplementary Data 3).

In *A. thaliana,* the feeding-induced genes were additionally significantly enriched in KEGG pathway terms related to glucosinolate biosynthesis [e.g., *SULFOTRANSFERASE 16 and 17* (*SOT16*, *SOT17*)], biosynthesis of various secondary metabolites and plant hormone signal transduction [*JASMONATE ZIM-DOMAIN PROTEIN 1* (*JAZ1*)] (Fig. 5, Supplementary Data 2).

#### Downregulation of genes in response to larval feeding

In both plant species, exposure to larval feeding downregulated genes associated with the KEGG pathway terms biosynthesis of secondary metabolites, fatty acid biosynthesis and metabolism, carbon metabolism, fixation in photosynthetic organisms, and porphyrin metabolism (Fig. 5). Additionally, the GO term analysis showed that in both plant species, photosynthesis-related genes were downregulated in response to larval feeding (Supplementary Data 3).

Overall, the downregulated genes in feeding-damaged *A. lyrata* were enriched in more KEGG pathway and GO terms than in *A. thaliana* [KEGG pathway term enrichment: 32 terms for *A. lyrata* and 17 terms for *A. thaliana* (Fig. 5, Supplementary Data 2); GO term enrichment: 121 terms for *A. lyrata* and 86 terms for *A. thaliana* (Supplementary Data 3)].

In feeding-damaged *A. lyrata,* overall more downregulated genes were significantly enriched in KEGG pathway and GO terms that are associated to photosynthesis and pigment synthesis (e.g., KEGG pathways: photosynthesis-antenna proteins, photosynthesis [(*PHOTOSYSTEM II REACTION CENTER PSB28 PROTEIN* (*PSB28*), *PHOTOSYSTEM I SUBUNIT D-2* (*PSAD-2*)], chlorophyll synthesis (*CHLD* encoding a subunit of the magnesium chelatase), carotenoid biosynthesis, and glucosinolate biosynthesis [e.g., *ISOPROPYL MALATE ISOMERASE LARGE SUBUNIT 1* (*IIL1*), *SOT17*] (Fig. 5, Supplementary Data 2).

### Responses to eggs exert negligible effects on larval feeding-damaged *Arabidopsis lyrata*

Finally, we compared the impact of *P. brassicae* eggs and *P. brassicae* larval feeding on the transcriptomes of seven-week-old *A. thaliana* and *A. lyrata* with RNA-seq.

In *A. thaliana*, 58% of egg-inducible genes (E vs C) were also induced in response to larval feeding (F vs C) (582 genes) (Fig. 6a). In *A. lyrata,* 77% of egg-responsive genes were also induced by larval feeding (274 commonly upregulated genes from 354 in total; Fig. 6a). However, compared to *A. thaliana*, *A. lyrata* upregulated less than 30% of the genes in response to eggs (Fig. 3, E vs C). From in total 5977 upregulated genes throughout all treatments, only 354 (less than 6%) were upregulated in response to eggs in *A. lyrata* (Fig. 6a).

**Fig. 6.**
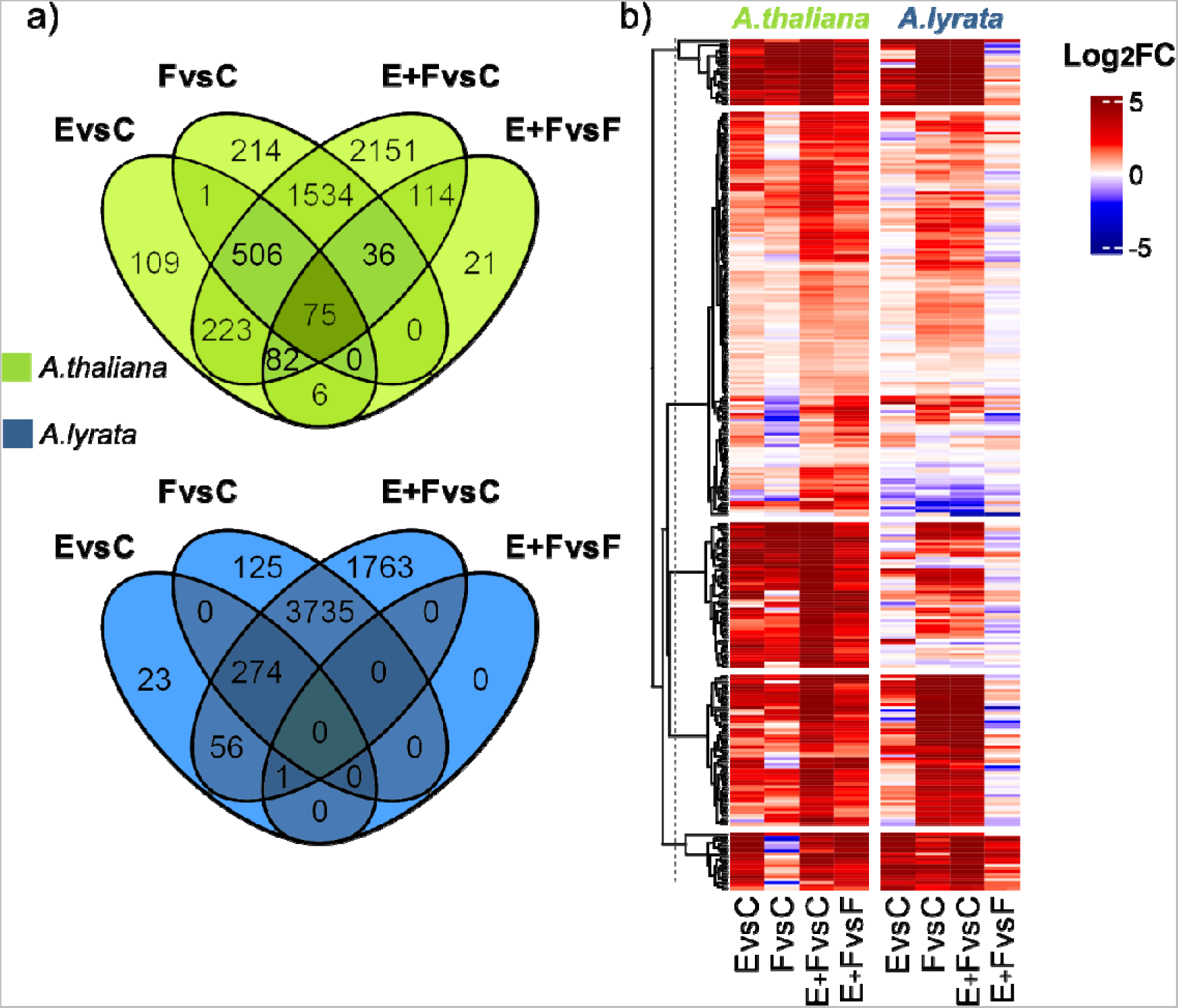
Impact of *Pieris brassicae* eggs and larval feeding to the primed transcriptome of *Arabidopsis thaliana* or *A. lyrata*. a) Venn diagrams indicate the number of upregulated genes in *A. thaliana* (green) and *A. lyrata* (blue) that are commonly or uniquely regulated in response to *P. brassicae* eggs (E), larval feeding (F) or eggs and larval feeding (E+F). C represents transcriptomes of untreated plants. b) Hierarchically clustered heatmap of up- or down-regulated *A. thaliana* and *A. lyrata* genes for the following treatment comparisons: E versus C, F versus C, E+F versus C and E+F versus F. Red colors indicate upregulation, blue colors downregulation of genes. *N* = 5

When comparing the upregulated DEGs in E+F plants versus those in F plants in *A. lyrata,* we identified one cluster of genes that tended to be upregulated (Fig. 6b). However, only one gene (AT2G45340, encoding a leucine-rich repeat protein kinase family protein) was significantly upregulated in the E+F plants compared to F plants (E+F vs F, Fig. 6a, Supplementary Data 1 and 5).

By contrast, *A. thaliana* upregulated 334 genes in E+F plants compared to only feeding-damaged F plants (Fig. 6a, Supplementary Data 1). Among those are *CAX3* and *PR1* (Fig. 4), which were identified as egg-primed genes in previous studies (Bruessow et al. 2010; Little et al. 2007; Lortzing et al. 2019; Valsamakis et al. 2022). These 334 upregulated genes play a role, for example, in biosynthesis of phenylpropanoids [e.g., several genes coding for peroxidases (*PER23*, *PER50*, *PER54*, *PER58*), *4CLL7* that encodes 4-coumarate-CoA ligase-like 7] and in immune responses (e.g., *WRKY41*, *WRKY47*) (Fig. 5, Supplementary Data 2 and 3).

Similar patterns were detected for downregulated genes when comparing E+F vs F plants (Fig. S2a). Whereas in *A. lyrata* only one gene was downregulated in E+F plants when compared to F plants (AT4G33550, encoding a bifunctional inhibitor/lipid-transfer protein/seed storage 2S albumin superfamily protein) (Supplementary Data 1), *A. thaliana* E+F plants downregulated 201 genes when compared to F plants (Fig. S2a). These 201 genes were involved, for example, in chlorophyll biosynthesis [e.g., *UROPORPHYRINOGEN III SYNTHASE* (*DUF3*)] and in photosynthesis [e.g., *PHOTOSYSTEM II SUBUNIT T* (*PSBTN*)] (Supplementary Data 3).

The small transcriptomic differences between egg-laden and feeding-damaged (E+F) *A. lyrata* plants and plants that were only fed upon (F), compared to the response of equally treated *A. thaliana* indicates the absence of an egg-mediated priming response in *A. lyrata* (Fig. 6b, Fig. S2b).

## Discussion

Our study showed that defense of the perennial Brassicaceae *A. lyrata* against larvae of *P. brassicae* is not primable by prior egg deposition. Thus, this perennial *A. lyrata* shows responses to *P. brassicae* eggs and larvae that differ from those known for other annual Brassicaceae and for the perennial *B. oleracea*. (Pashalidou et al. 2013; Pashalidou et al. 2015; Pashalidou et al. 2020). Here, we specifically compared the egg-primability of the anti-herbivore defense of the perennial *A. lyrata* with the one of the closely related annual *A. thaliana*.

As in previous studies (Geiselhardt et al. 2013; Lortzing et al. 2019; Paniagua Voirol et al. 2020; Valsamakis et al. 2020; Valsamakis et al. 2022), *P. brassicae* larvae gained less biomass on previously egg-laden *A. thaliana* plants compared to larvae on egg-free plants. By contrast, this priming effect was absent in *A. lyrata*, i.e., the larvae performed equally well on previously egg-laden and egg-free plants (Fig. 1). The absence of the priming effect in *A. lyrata* is also reflected by its transcriptomic response to *P. brassicae* eggs with subsequent larval feeding. Our RNA-seq analysis revealed that only one gene, *AT2G45340* encoding for a leucine-rich repeat protein kinase family protein, was stronger induced in feeding-damaged *A. lyrata* when plants had previously received the eggs (Fig. 2 and Fig. 6a, Supplementary Data 1). In contrast, prior egg deposition on feeding-damaged *A. thaliana* resulted in stronger induction of 334 genes compared to feeding-damaged plants without prior egg deposition (Fig. 2 and Fig. 6a), including typically egg priming-responsive genes like *PR1*, *PR5* and *CAX3* and several *WRKY* transcription factor genes (Fig. 4, Supplementary Data 1). The 334 upregulated priming-responsive genes in *A. thaliana* play amongst others a role in phenylpropanoid biosynthesis and immune responses, including responses to SA (Fig. 5, Supplementary Data 2 and 3), which was shown to be crucial for establishing the egg-mediated anti-herbivore defense response in *A. thaliana* (Lortzing et al. 2019; Valsamakis et al. 2020).

*Arabidopsis lyrata* upregulated considerably fewer genes in response to eggs than *A. thaliana*. The rather moderate transcriptional response of *A. lyrata* to *P. brassicae* eggs might result in the absence of a priming response in *A. lyrata.* Although *A. lyrata* genes were responsive to eggs *per se*, this transcriptional change did not result in an additive or synergistic transcriptional interactions with the “larval feeding stimulus” (Fig. 6b). By contrast, *A. thaliana* responded much stronger to the *P. brassicae* eggs resulting in maintained egg-induced responses or additive and synergistic interactions between “egg stimulus” and “larval feeding stimulus” (Fig. 6b). Additive and synergistic effects contribute to the priming response of *A. thaliana* and result in an accelerated and stronger resistance response of *A. thaliana* against *P. brassicae* larvae if the plant is previously exposed to conspecific eggs (Lortzing et al. 2020; Valsamakis et al. 2020; Valsamakis et al. 2022).

The expression of genes typically responsive to eggs, such as *PDF1.4, PR1*, and *PR5* (Little et al. 2007; Paniagua Voirol et al. 2020; Valsamakis et al. 2020) was significantly induced by eggs alone in both plant species (Fig. 4). By contrast, the expression of *CAX3* was strongly induced by eggs in *A. thaliana,* but not in *A. lyrata* (Fig. 4). *CAX3* encodes for a Ca^2+^/H^+^ exchanger localized in the tonoplast (Manohar et al. 2011). Thus, calcium signaling seems to play an important role in perceiving the *P. brassicae* eggs and establishing the egg-primed plant defense response against the larvae.

Furthermore, prior egg deposition on feeding-damaged *A. thaliana* resulted in upregulation of genes that are involved in oxidative stress (Supplementary Data 3). Calcium signaling and oxidative stress are linked (Ermak and Davies 2002). Oxidative stress occurs when the balance between reactive oxygen species (ROS) and antioxidant molecules is disrupted (Scandalios 2002) and Ca^2+^ peaks in the cytoplasm are induced (Ermak and Davies 2002). When applied onto plant wounds, insect-derived elicitors, e.g., inceptin and volicitin, induce a Ca^2+^ influx, which in turn triggers ROS production (Kumar et al. 2020; Schmelz et al. 2006). During stress such as herbivory, the level of cytosolic Ca^2+^ rapidly increases due to Ca^2+^ influx mediated by Ca^2+^-ATPases and Ca^2+^ efflux mediated by H^+^/ Ca^2+^ exchanger such as CAX1 and CAX3. CAX1 and CAX3 have high specificity for Ca^2+^ binding and maintenance of Ca^2+^ homeostasis (Kumar et al. 2020). Furthermore, ROS is known to locally accumulate at the oviposition site in various annual and perennial plant species (Bittner et al. 2017; Geuss et al. 2017; Gouhier-Darimont et al. 2013; Little et al. 2007; Reymond 2013). ROS accumulation and cytosolic Ca^2+^ increase might initiate *A. thaliana*’s response to eggs and contribute to mounting defenses against *P. brassicae* larvae. Further investigation will shed light to the yet unknown role of CAXs and on how oxidative stress and calcium signaling are integrated to establish the egg-primed anti-herbivore defense in *A. thaliana*.

Although *A. lyrata* responds to *P. brassicae* eggs only with moderate transcriptional changes, its transcriptional response to larval feeding without prior egg deposition was much stronger than that of *A. thaliana* (Fig. 2, Fig. 6a, Supplementary Data 1). In response to larval feeding, both plant species upregulated genes that play a role in JA and SA synthesis and signaling as well as in the synthesis of flavonoids including anthocyanins. Furthermore, both plant species downregulated genes that play a role in photosynthesis (Fig. 5, Supplementary Data 2 and 3). Previous studies showed that plant defense responses to insect herbivore attack are associated with a reduction in photosynthesis, thus allowing plants to allocate resources toward immediate defense needs (Bilgin et al. 2010; Kerchev et al. 2012). However, regarding upregulated genes that are involved in SA-mediated signaling and downregulated genes that are involved in photosynthesis, *A. lyrata*’s response to larval feeding seems to be more pronounced than *A. thaliana*’s response. The stronger feeding-mediated induction of SA-responsive and defense-related genes was also reflected by the stronger expression of *PR1*, *PR5* and *PDF.1.4* in *A. lyrata* than in *A. thaliana* (Fig. 4). Additionally, *CAX3* expression was remarkably strong in response to larval feeding (Fig. 4). Therefore, Ca^2+^ signaling might be a key component for *A. lyrata’*s defense response against the *P. brassicae* larvae. A study from Toyota et al. (2018) revealed that feeding by *P. rapae* larvae induced cytosolic Ca^2+^ accumulation in *A. thaliana*. Thus, *P. brassicae* larval feeding may affect *A. lyrata* in a similar manner as *P. rapae* feeding on *A. thaliana*.

Taken together, the massive transcriptome reprogramming of *A. lyrata* in response to larval feeding might compensate for *A. lyrata*’s weak responsiveness to *P. brassicae* eggs and its lack of egg-mediated primability.

It is tempting to attribute the defense strategies of *A. thaliana* and *A. lyrata* to their respective natural habitats. Since priming against herbivory is costly (Valsamakis et al. 2022), growing in environments with low density of competing species (Mitchell-Olds 2001; Vergeer and Kunin 2011) may push *A. lyrata* to prioritize its stress responses, thus possibly saving resources for growth (Saijo and Loo 2020; Wise and Abrahamson 2007). If stress by herbivory occurs relative to other stress events less frequently in the habitat of *A. lyrata,* it might be cost-saving when limiting primability to the most frequent stress responses. If this is the case with *A. lyrata* habitats, it might explain why *A. lyrata* remains unaffected by *P. brassicae* eggs and exhibits a stronger response once larvae start feeding, to compensate for the lack of egg-mediated priming. Conversely, given that *A. thaliana* prospers mostly in environments where nutrients and water are not limited (Al-Shehbaz and O’Kane 2002) and where herbivory might represent a more frequently occurring threat, for *A. thaliana* the more efficient strategy would be to invest in defense against herbivory after egg deposition, that is, by priming its defenses before the larvae hatch. Other egg-primable Brassicaceae species like *B. nigra* (Pashalidou et al. 2013), *B. oleracea* and *S. arvensis* (Pashalidou et al. 2015) also occur on soils where nutrient and water are usually not limited (Fogg 1950; Rich 1991; Stace 1997).

In conclusion, our results suggest that the primability of anti-herbivore defenses against herbivory is not phylogenetically conserved in the genus *Arabidopsis*. In contrast to other perennial plants – including *B. oleracea* as a Brassicaceae species – *A. lyrata’s* anti-herbivore defense turned out to be not primable by insect egg deposition, indicating that the life span of a plant does not affect its egg primability (Beyaert et al. 2012; Austel et al. 2016; Pashalidou et al. 2015a; Geuss et al. 2018). Future studies are needed to elucidate the traits that determine a plant’s sensitivity to insect eggs, which probably affect the plant’s egg primability. The balance between the frequency of insect infestations and other (a)biotic stresses in the natural habitats of plants might shape the selection for traits relevant for becoming primed by insect egg deposition. Among these traits, those responsive for perception of eggs and the specific role of *CAX3* in plant defense responses against herbivores needs to be addressed in future studies.

## Data accessibility

All presented data are included in the article or are available as Supplementary data. The RNA-seq raw sequences data are deposited at the European Bioinformatics Institute (EBI) platforms ArrayExpress and Expression Atlas under the accession number E-MTAB-12653.

## Author contribution statement

NB, RK, MH, MRE, LPV and VL conceptualized the study. NB, MHu and VL performed and analyzed the experiments. MHu and NB wrote the first draft. MH, RK, MRE, LPV and VL contributed to later versions of the manuscript and agreed with the final version.

## Supporting information

List of differentially expressed genes

KEGG pathway term enrichment

GO term enrichment

List of DEG between species

List of DEG between treatments

## Acknowledgement

Many thanks to Dr. Marc Stift from the University of Konstanz, Germany, for providing the *A. lyrata* selfing-line. We are grateful to Laura Hagemann, Freie Universität Berlin (FUB), Germany, for helping with the insect and plant rearing and assistance during the experiments. Thanks to the High-Performance Computing cluster (FUB) for providing computing time. We thank the German Research Foundation (DFG) for funding (DFG project 502563004 and Collaborative Research Centre 973, project B4: https://www.sfb973.de).

## Main conclusion

Unlike *Arabidopsis thaliana*, defenses of *Arabidopsis lyrata* against *Pieris brassicae* larval feeding are not primable by *P. brassicae* eggs. Therefore, egg primability of plant anti-herbivore defenses is not phylogenetically conserved in the genus *Arabidopsis*.

## Supporting information

### Supplementary Data

**Supplementary Data 1** List of differentially expressed genes (upregulated: red; downregulated: blue) in *Arabidopsis thaliana* and *A. lyrata* in response to *Pieris brassicae* eggs (E vs. C), larval feeding (F vs. C) or both, eggs and larval feeding (E+F vs. C; E+F vs. F). *N* = 5

**Supplementary Data 2** KEGG pathway term enrichment of differentially expressed genes in *Arabidopsis thaliana* and *A. lyrata:* All treatment comparisons. Plants were exposed to *Pieris brassicae* eggs (E), larval feeding (F), or eggs and subsequent larval feeding (E+F) compared to untreated controls (C) or feeding-damaged plants with prior egg deposition compared to feeding-damaged plants without prior egg deposition (E+F vs F). The tables show KEGG term enrichment for upregulated (up) and downregulated (down) genes. *N* = 5

**Supplementary Data 3** Biological process GO term enrichment of differentially expressed genes in *Arabidopsis thaliana* and *A. lyrata:* All treatment comparisons. Plants were exposed to *Pieris brassicae* eggs (E), larval feeding (F), or eggs and subsequent larval feeding (E+F) compared to untreated controls (C) or feeding-damaged plants with prior egg deposition compared to feeding-damaged plants without prior egg deposition (E+F vs F). The tables show GO term enrichment for upregulated (up) and downregulated (down) genes. *N* = 5

**Supplementary Data 4** List of differentially expressed genes, which are commonly or exclusively up- or downregulated in *Arabidopsis thaliana* (*AT*) and *A. lyrata* (*AL*) in response to *Pieris brassicae* eggs (E vs. C) or larval feeding (F vs. C). *N* = 5

**Supplementary Data 5** List of differentially commonly and exclusively expressed genes within *Arabidopsis thaliana* (*AT*) or *A. lyrata* (*AL*) in response to the treatments: *Pieris brassicae* eggs (E), larval feeding (F) or both (E+F). C plants were left untreated. *N* = 5

### Supplementary Figures

**Fig. S1** Impact of the plant’s responses to *Pieris brassicae* eggs on biomass of conspecific larvae feeding on nine-week-old *Arabidopsis lyrata* plants. Biomass in mg (means ± SE) of larvae after 2 or 5 days feeding on previously egg-laden (E+F) and egg-free (F) plants. Dots represent the data points. ns above the bars indicate non-significant differences between the treatments (*P* > 0.05, multiple Student’s *t*-test with *fdr* correction *post hoc*). *N* = 9

**Fig. S2** Impact of *Pieris brassicae* eggs and larval feeding to the primed transcriptome of *Arabidopsis thaliana* or *A. lyrata*. a) Venn diagrams indicate the number of downregulated genes in *A. thaliana* (green) and *A. lyrata* (blue) that are commonly or uniquely regulated in response to *P. brassicae* eggs (E), larval feeding (F) or eggs and larval feeding (E+F). C represents transcriptomes of untreated plants. b) Hierarchically clustered heatmap of up- or downregulated *A. thaliana* and *A. lyrata* genes for the following treatment comparisons: E versus C, F versus C, E+F versus C and E+F versus F. Red colors indicate upregulation, blue colors downregulation of genes. *N* = 5

### Supplementary Tables

**Supplementary Table S1** Sequences of primers used for qPCR

**Supplementary Table S2** Statistical details of data presented in Fig. 1

**Supplementary Table S3** Statistical details of data presented in Fig. 4

