## Supplemental Information for "Butterfly eggs prime anti-herbivore defense in an annual but not perennial *Arabidopsis* species"

^1^ Microbiology, Institute of Biology, Dahlem Centre of Plant Sciences, Freie Universität Berlin, Königin-Luise-Str. 12-16, 14195 Berlin, Germany; ^2^ Applied Genetics, Institute of Biology, Dahlem Centre of Plant Sciences, Freie Universität Berlin, Albrecht-Thaer-Weg 6, 14195 Berlin, Germany; ^3^ Applied Zoology / Animal Ecology, Institute of Biology, Dahlem Centre of Plant Sciences, Freie Universität Berlin, Haderslebener Str. 9, 12163 Berlin, Germany

* Correspondence: Vivien Lortzing (0000-0002-1480-0274) and Luis R. Paniagua Voirol (0000-0001-9868-1884)

^a^ These authors contributed equally to this work (MHu, NB)

**Overview**

**Supplementary Data in this file**

**Fig. S1** Impact of the plant’s responses to *Pieris brassicae* eggs on biomass of conspecific larvae feeding on nine-week-old *Arabidopsis lyrata* plants. Biomass in mg (means ± SE) of larvae after 2 or 5 days feeding on previously egg-laden (E+F) and egg-free (F) plants. Dots represent the data points. ns above the bars indicate non-significant differences between the treatments (*P* > 0.05, multiple Student’s *t*-test with *fdr* correction *post hoc*). *N* = 9

**Fig. S2** Impact of *Pieris brassicae* eggs and larval feeding to the primed transcriptome of *Arabidopsis thaliana* or *A. lyrata*. a) Venn diagrams indicate the number of downregulated genes in *A. thaliana* (green) and *A. lyrata* (blue) that are commonly or uniquely regulated in response to *P. brassicae* eggs (E), larval feeding (F) or eggs and larval feeding (E+F). C represents transcriptomes of untreated plants. b) Hierarchically clustered heatmap of up- or downregulated *A. thaliana* and *A. lyrata* genes for the following treatment comparisons: E versus C, F versus C, E+F versus C and E+F versus F. Red colors indicate upregulation, blue colors downregulation of genes. *N* = 5

**Supplementary Table S1** Sequences of primers used for qPCR

**Supplementary Table S2** Statistical details of data presented in Fig. 1

**Supplementary Table S3** Statistical details of data presented in Fig. 4

**Supplementary Data available in additional ZIP*:***

**Supplementary Data 1** List of differentially expressed genes (upregulated: red; downregulated: blue) in *Arabidopsis thaliana* and *A. lyrata* in response to *Pieris brassicae* eggs (E vs. C), larval feeding (F vs. C) or both, eggs and larval feeding (E+F vs. C; E+F vs. F). *N* = 5

**Supplementary Data 2** KEGG pathway term enrichment of differentially expressed genes in *Arabidopsis thaliana* and *A. lyrata:* All treatment comparisons. Plants were exposed to *Pieris brassicae* eggs (E), larval feeding (F), or eggs and subsequent larval feeding (E+F) compared to untreated controls (C) or feeding-damaged plants with prior egg deposition compared to feeding-damaged plants without prior egg deposition (E+F vs F). The tables show KEGG term enrichment for upregulated (up) and downregulated (down) genes. *N* = 5

**Supplementary Data 3** Biological process GO term enrichment of differentially expressed genes in *Arabidopsis thaliana* and *A. lyrata:* All treatment comparisons. Plants were exposed to *Pieris brassicae* eggs (E), larval feeding (F), or eggs and subsequent larval feeding (E+F) compared to untreated controls (C) or feeding-damaged plants with prior egg deposition compared to feeding-damaged plants without prior egg deposition (E+F vs F). The tables show GO term enrichment for upregulated (up) and downregulated (down) genes. *N* = 5

**Supplementary Data 4** List of differentially expressed genes, which are commonly or exclusively up- or downregulated in *Arabidopsis thaliana* (*AT*) and *A. lyrata* (*AL*) in response to *Pieris brassicae* eggs (E vs. C) or larval feeding (F vs. C). *N* = 5

**Supplementary Data 5** List of differentially commonly and exclusively expressed genes within *Arabidopsis thaliana* (*AT*) or *A. lyrata* (*AL*) in response to the treatments: *Pieris brassicae* eggs (E), larval feeding (F) or both (E+F). C plants were left untreated. *N* = 5

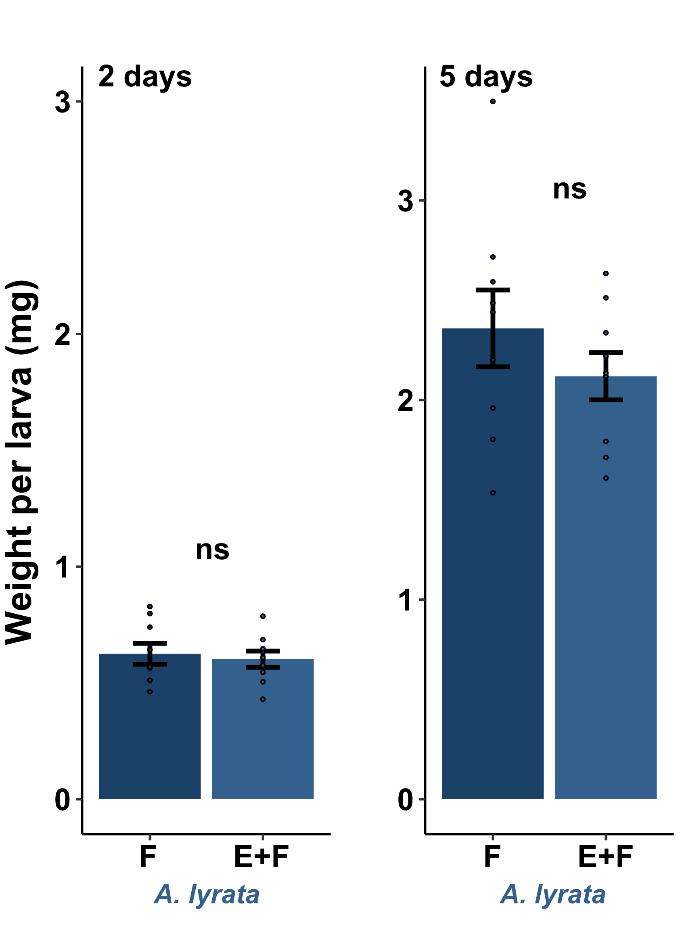

**Fig. S1** Impact of the plant’s responses to *Pieris brassicae* eggs on biomass of conspecific larvae feeding on nine-week-old *Arabidopsis lyrata* plants. Biomass in mg (means ± SE) of larvae after 2 or 5 days feeding on previously egg-laden (E+F) and egg-free (F) plants. Dots represent the data points. ns above the bars indicate non-significant differences between the treatments (*P* > 0.05, multiple Student’s *t*-test with *fdr* correction *post hoc*). *N* = 9

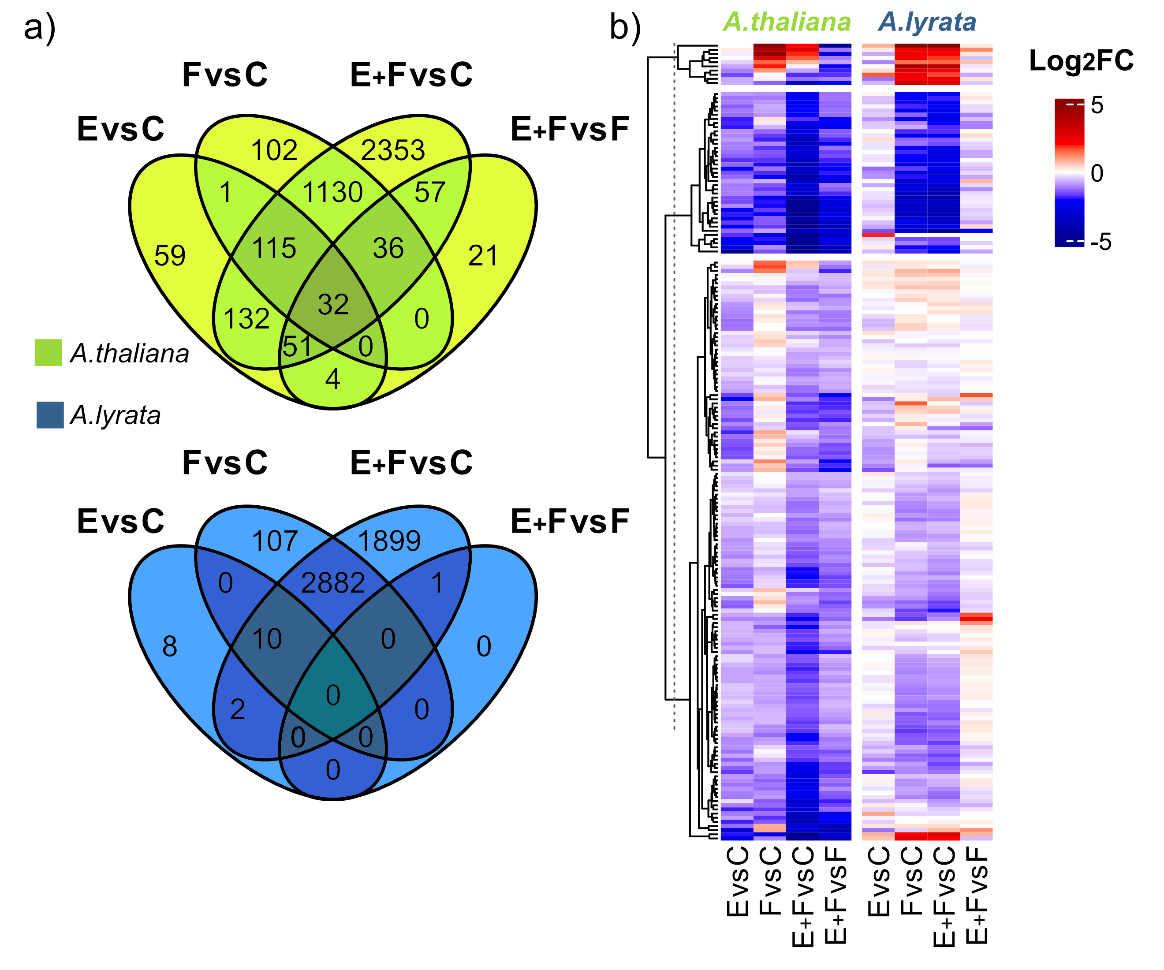

**Fig. S2** Impact of *Pieris brassicae* eggs and larval feeding to the primed transcriptome of *Arabidopsis thaliana* or *A. lyrata*. a) Venn diagrams indicate the number of downregulated genes in *A. thaliana* (green) and *A. lyrata* (blue) that are commonly or uniquely regulated in response to *P. brassicae* eggs (E), larval feeding (F) or eggs and larval feeding (E+F). C represents transcriptomes of untreated plants. b) Hierarchically clustered heatmap of up- or downregulated *A. thaliana* and *A. lyrata* genes for the following treatment comparisons: E versus C, F versus C, E+F versus C and E+F versus F. Red colors indicate upregulation, blue colors downregulation of genes. *N* = 5

**Supplementary Table S1** Sequences of primers used for qPCR

|  |  | **Sequence 5´→3´** | |
| --- | --- | --- | --- |
| **Target** |  | **Forward** | **Reverse** |
| ***Arabidopsis thaliana*** | | | |
| ***ACT2*** | AT3G18780 | CTTCCCTCAGCACATTCCAG | GACCTGCCTCATCATACTCG |
| ***TUB6*** | AT5G12250 | ACCACTCCTAGCTTTGGTGATCTG | AGGTTCACTGCGAGCTTCCTCA |
| ***GAPDH*** | AT1G13440 | TTGGTGACAACAGGTCAAGCA | AAACTTGTCGCTCAATGCAATC |
| ***PR1*** | AT2G14610 | ACTTAGCCTGGGGTAGCGGT | CCATTGCACGTGTTCGCAGC |
| ***PR5*** | AT1G75040 | ATCACCCACAGCACAGAGACAC | AGCAATGCCGCTTGTGATGAAC |
| ***CAX3*** | AT3G51860 | TTTGTTCGTGGTCCCATTGAG | CATCCTGTAAAGTGAAGGCTGTG |
| ***PDF1.4*** | AT1G19610 | TGTGTGAATCTTCTTCAACGGTCAC | AGAAGATAGCCTTGCCAGTCTCTTA |
| ***Arabidopsis lyrata*** | | | |
| ***ALPS2SUB*** | XM_002887719 | AGCTAACGTCGATGGATACAGTCC | AAATTGCCAACCCGGTGACTCC |
| ***ALPYRT*** | XM_021030662 | CCGGAGCAGAGATTACACAAATCC | AAGCAGGCTGCATAGACCCATC |
| ***ALPPP2R1* 3'** | XM_002892696 | AGACAAGGTTCACTCAATCCGTG | CATTCAGGACCAAACTCTTCAGC |
| ***ALCAX3*** | XM_021025964 | CGCCATCCCTCTCGCCATTC | AGCTCCGTTGCGTTTCCACA |
| ***AL_PDF1.4*** | XM_021014212 | TTGCGAGAGGCGAAGCAAGA | CGTGACAAGCCCCGTGTGAA |
| ***ALPR1*** | XM_002885879 | TTTGCGGACACTACACTCAGGTC | ACTTTAGCACATCCGAGTCTCACC |
| ***AL PR5*** | XM_002887525 | TCTGCCCATAAAAAGGCGAAGC | ACAAGCAAAAGAAACGTGAGAGGT |

**Supplementary Table S2** Statistical details of data presented in Fig. 1

**Experiment 1:** Statistical differences (*P values)* between biomass of larva after 2 days feeding on egg-laden (E+F) or egg-free (F) *Arabidopsis* *thaliana* (*AT*) or *A.* *lyrata* (*AL*) plants (multiple Student’s *t*-tests with *fdr* adjusted *P* values).

| ***P*** | *AL*-E+F | *AL*-F | *AT*-E+F | ***N*** | F | E+F |
| --- | --- | --- | --- | --- | --- | --- |
| *AL*-F | 0.80793 | - | - | *AL* | 17 | 16 |
| *AT*-E+F | 0.00014 | 0.00016 | - | *AT* | 18 | 18 |
| *AT*-F | 0.01298 | 0.02268 | 0.01147 |  |  |  |

**Experiment 2:** Statistical differences (*P values)* between biomass of larva after 5 days feeding on egg-laden (E+F) or egg-free (F) *Arabidopsis* *thaliana* (*AT*) or *A.* *lyrata* (*AL*) plants (multiple Student’s *t*-tests with *fdr* adjusted *P* values).

| ***P*** | *AL*-E+F | *AL*-F | *AT*-E+F | ***N*** | F | E+F |
| --- | --- | --- | --- | --- | --- | --- |
| *AL*-F | 0.15085 | - | - | *AL* | 8 | 7 |
| *AT*-E+F | 1.3e^-06^ | 3.3e^-08^ | - | *AT* | 8 | 8 |
| *AT*-F | 0.00096 | 1.5e^-05^ | 0.01003 |  |  |  |

**Supplementary Table S3** Statistical details of data presented in Fig. 4

| ***Arabidopsis thaliana*** | ***Arabidopsis lyrata*** |
| --- | --- |
| ***CAX3*** |  |
| ***2x2 ANOVA*** |  |
| \| Df \| Sum Sq \| Mean Sq \| F value \| Pr(>F); E \| 1 \| 205.65 \| 205.65 \| 51.05 \| 2.33E^-06^; F \| 1 \| 24.41 \| 24.41 \| 6.058 \| 0.0256; E:F \| 1 \| 17.6 \| 17.6 \| 4.368 \| 0.0529; Residuals \| 16 \| 64.46 \| 4.03 \| \| | \| Df \| Sum Sq \| Mean Sq \| F value \| Pr(>F); E \| 1 \| 205.65 \| 205.65 \| 51.05 \| 2.33E^-06^; F \| 1 \| 24.41 \| 24.41 \| 6.058 \| 0.0256; E:F \| 1 \| 17.6 \| 17.6 \| 4.368 \| 0.0529; Residuals \| 16 \| 64.46 \| 4.03 \| \| |
| **Tukey multiple comparisons of means** |  |
| \| diff \| lwr \| upr \| p adj; F:E+F \| -4.537333 \| -8.169135 \| -0.905532 \| 0.012169; E:E+F \| -0.333333 \| -3.965135 \| 3.298468 \| 0.993402; C:E+F \| -8.622667 \| -12.25447 \| -4.990865 \| 0.000023; E:F \| 4.204 \| 0.572198 \| 7.835802 \| 0.020642; C:F \| -4.085333 \| -7.717135 \| -0.453532 \| 0.024870; C:E \| -8.289333 \| -11.92114 \| -4.657532 \| 0.000037 | \| diff \| lwr \| upr \| p adj; F:E+F \| 1.528667 \| -1.489249 \| 4.5465826 \| 0.4888862; E:E+F \| -2.649333 \| -5.667249 \| 0.3685826 \| 0.0960127; C:E+F \| -3.905333 \| -6.923249 \| -0.8874174 \| 0.0093886; E:F \| -4.178 \| -7.195916 \| -1.1600841 \| 0.0055488; C:F \| -5.434 \| -8.451916 \| -2.4160841 \| 0.0005054; C:E \| -1.256 \| -4.273916 \| 1.7619159 \| 0.6411177 |
| ***PDF1.4*** |  |
| ***2x2 ANOVA*** |  |
| \| Df \| Sum Sq \| Mean Sq \| F value \| Pr(>F); E \| 1 \| 50.13 \| 50.13 \| 33.966 \| 2.57E^-05^; F \| 1 \| 3.6 \| 3.6 \| 2.438 \| 0.13798; E:F \| 1 \| 16.8 \| 16.8 \| 11.383 \| 0.00387; Residuals \| 16 \| 23.61 \| 1.48 \| \| | \| Df \| Sum Sq \| Mean Sq \| F value \| Pr(>F); E \| 1 \| 87.52 \| 87.52 \| 25.59 \| 0.000116; F \| 1 \| 130.34 \| 130.34 \| 38.12 \| 1.34E^-05^; E:F \| 1 \| 38.26 \| 38.26 \| 11.19 \| 0.00411; Residuals \| 16 \| 54.71 \| 3.42 \| \| |
| **Tukey multiple comparisons of means** |  |
| \| diff \| lwr \| upr \| p adj; F:E+F \| -1.333333 \| -3.531561 \| 0.864895 \| 0.338756; E:E+F \| 0.984667 \| -1.213561 \| 3.182895 \| 0.58697; C:E+F \| -4.014667 \| -6.212894 \| -1.816439 \| 0.000437; E:F \| 2.318 \| 0.119772 \| 4.516228 \| 0.03698; C:F \| -2.681333 \| -4.879561 \| -0.483105 \| 0.014437; C:E \| -4.999333 \| -7.197561 \| -2.801105 \| 0.014437 | \| diff \| lwr \| upr \| p adj; F:E+F \| -1.417333 \| -4.763421 \| 1.928755 \| 0.6285157; E:E+F \| -2.339333 \| -5.685421 \| 1.006755 \| 0.2287714; C:E+F \| -9.289333 \| -12.63542 \| -5.943245 \| 0.0000033; E:F \| -0.922 \| -4.268088 \| 2.424088 \| 0.8586633; C:F \| -7.872 \| -11.21809 \| -4.525912 \| 0.0000261; C:E \| -6.95 \| -10.29609 \| -3.603912 \| 0.0001103 |
| ***PR1*** |  |
| ***2x2 ANOVA*** |  |
| \| Df \| Sum Sq \| Mean Sq \| F value \| Pr(>F); E \| 1 \| 217.36 \| 217.36 \| 39.484 \| 1.09E^-05^; F \| 1 \| 0.57 \| 0.57 \| 0.104 \| 0.751; E:F \| 1 \| 7.42 \| 7.42 \| 1.347 \| 0.263; Residuals \| 16 \| 88.08 \| 5.51 \| \| | \| Df \| Sum Sq \| Mean Sq \| F value \| Pr(>F); E \| 1 \| 108.1 \| 108.1 \| 21.426 \| 0.000279; F \| 1 \| 22.93 \| 22.93 \| 4.546 \| 0.048849; E:F \| 1 \| 50.07 \| 50.07 \| 9.923 \| 0.006194; Residuals \| 16 \| 80.72 \| 5.05 \| \| |
| **Tukey multiple comparisons of means** |  |
| \| diff \| lwr \| upr \| p adj; F:E+F \| -5.375333 \| -9.62087 \| -1.12980 \| 0.011041; E:E+F \| 1.556667 \| -2.68887 \| 5.80221 \| 0.723991; C:E+F \| -6.254667 \| -10.50021 \| -2.00913 \| 0.003308; E:F \| 6.932 \| 2.68646 \| 11.17754 \| 0.001314; C:F \| -0.879333 \| -5.12487 \| 3.36621 \| 0.932866; C:E \| -7.811333 \| -12.05687 \| -3.56580 \| 0.000405 | \| diff \| lwr \| upr \| p adj; F:E+F \| -1.485333 \| -5.549675 \| 2.579008 \| 0.7259543; E:E+F \| 1.022667 \| -3.041675 \| 5.087008 \| 0.8877362; C:E+F \| -6.791333 \| -10.85568 \| -2.726992 \| 0.0010553; E:F \| 2.508 \| -1.556342 \| 6.572342 \| 0.324743; C:F \| -5.306 \| -9.370342 \| -1.241658 \| 0.0087842; C:E \| -7.814 \| -11.87834 \| -3.749658 \| 0.000256 |
| ***PR5*** |  |
| ***2x2 ANOVA*** |  |
| \| Df \| Sum Sq \| Mean Sq \| F value \| Pr(>F); E \| 1 \| 41.91 \| 41.91 \| 25.153 \| 0.000127; F \| 1 \| 0.16 \| 0.16 \| 0.099 \| 0.757502; E:F \| 1 \| 11.74 \| 11.74 \| 7.042 \| 0.017338; Residuals \| 16 \| 26.66 \| 1.67 \| \| | \| Df \| Sum Sq \| Mean Sq \| F value \| Pr(>F); E \| 1 \| 45.68 \| 45.68 \| 40.31 \| 9.66E^-06^; F \| 1 \| 59.88 \| 59.88 \| 52.84 \| 1.88E^-06^; E:F \| 1 \| 25.48 \| 25.48 \| 22.48 \| 0.000221; Residuals \| 16 \| 18.13 \| 1.13 \| \| |
| **Tukey multiple comparisons of means** |  |
| \| diff \| lwr \| upr \| p adj; F:E+F \| -1.363333 \| -3.699175 \| 0.972508 \| 0.370511; E:E+F \| 1.350667 \| -0.985175 \| 3.686508 \| 0.378282; C:E+F \| -3.076667 \| -5.412508 \| -0.740825 \| 0.008209; E:F \| 2.714 \| 0.378157 \| 5.049841 \| 0.020136; C:F \| -1.713333 \| -4.049175 \| 0.622508 \| 0.195567; C:E \| -4.427333 \| -6.763175 \| -2.091492 \| 0.000298 | \| diff \| lwr \| upr \| p adj; F:E+F \| -0.765333 \| -2.691596 \| 1.1609295 \| 0.6730819; E:E+F \| -1.203333 \| -3.129596 \| 0.7229295 \| 0.3148285; C:E+F \| -6.483333 \| -8.409596 \| -4.557071 \| 0.0000003; E:F \| -0.438 \| -2.364263 \| 1.4882628 \| 0.91382; C:F \| -5.718 \| -7.644263 \| -3.791737 \| 0.0000014; C:E \| -5.28 \| -7.206263 \| -3.353737 \| 0.0000039 |
